# Androgenic Steroids Induce Pathologic Scarring in a Preclinical Swine Model Via Dysfunctional Extracellular Matrix Deposition

**DOI:** 10.1101/2023.05.29.542765

**Authors:** Erik Reiche, Patrick R Keller, Vance Soares, Calvin R Schuster, Siti Rahmayanti, Jessica Mroueh, Vanessa Mroueh, Marie Billaud, Sophia Hu, Hunter Hoover-Watson, Christine G Lian, Yu Tan, Joshua C Doloff, Annie E Newell-Fugate, Devin Coon

## Abstract

**Background:** Hypertrophic scarring is a major source of morbidity for surgery patients. Sex hormones are not classically considered to be modulators of scarring. However, based on clinical observations of increased frequency of hypertrophic scarring in patients on testosterone, we hypothesized that androgenic steroids induce abnormal scarring and developed a preclinical swine model to explore these effects.

**Methods:** A total of six male (XY) and female (XX) mini-swine underwent castration and were randomly assigned to no testosterone (noT) or biweekly testosterone therapy (+T). Ten dorsal excisional wounds were created on each pig. To mimic a chronic wound, a subset of wounds were re-excised at two weeks. Scars (POD42) and chronic wounds (POD28) were harvested six weeks after initial wounding for analysis via histology, RNA-seq, and mechanical testing.

**Results:** Histologic analysis of POD42 scars from +T swine showed increased mean fibrosis area (16mm^2^ noT, 28mm^2^ +T; p=0.007) and thickness (0.246mm^2^ noT, 0.406mm^2^ +T; p<0.001) compared to noT swine. Scars in XX+T and XY+T pigs had greater tensile burst strength (p=0.024 and p=0.013 respectively) compared to scars in noT swine. Color deconvolution analysis showed greater deposition of type I and type III collagen as well as increased type I to type III collagen ratio in +T scars. Dermatopathologist scores of POD42 scars show +T exposure was associated with worse overall scarring scores compared to controls (p<0.05). On RNAseq, gene ontology analysis showed testosterone exposure was associated with significant upregulation of cellular metabolism and immune response gene sets. Pathway analysis showed testosterone upregulated Reactome pathways related to keratinization and formation of collagen and laminin.

**Conclusion:** We developed a novel preclinical porcine model to study the effects of the sex hormone testosterone on scarring. Testosterone induces early proliferation of excessive granulation tissue, which eventually leads to increased scar tissue. T also appears to increase the physical strength of scars via supraphysiologic deposition of collagen and other ECM factors. The increase in burst strength observed for both XX and XY suggests that hormonal administration has a stronger influence on mechanical properties than chromosomal sex. Antiandrogen topical therapies may be a promising future area of research.

## Introduction

Hypertrophic scars with deposition of excessive scar tissue can affect up to 90% of burn or surgical patients and cause mental and physical issues, including pain and limited mobility, impairing quality of life.^1,2^ Scarring is a major source of postsurgical morbidity, representing both a physical and economic burden for patients. An estimated 100 million patients annually acquire scars from surgery, which can result in psychological distress and complications such as pain.^3–5^ Keloid and hypertrophic scarring in particular are associated with greater morbidity. In hypertrophic scars, myofibroblasts persist after epithelialization, leading to painful and functionally limiting skin contractures. Financially, there is an estimated $12 billion market in the United States for scar treatment.^6^

There are nearly 25 million transgender and gender-diverse (TGD) individuals worldwide receiving care that may include hormone replacement therapy and gender-affirming surgery (GAS).^7–10^ An increasing number of people are seeking GAS annually, making it urgent to address gaps in medical knowledge. Many patients receive exogenous testosterone therapy (T), and wound healing and scarring complications are a major source of morbidity for GAS patients. Despite this, while work has been published about the impact of testosterone on wound healing in murine models little is currently known about the impact of exogenous testosterone on scar formation. ^11–14^

In classic models of scarring, sex hormones (testosterone and estradiol) are not modulators of this process. However, over years of surgeries on patients on exogenous T, we observed slower WH and worse scarring, prompting our hypothesis that hormones were significant modulators of wound repair. We previously conducted a blinded analysis of scar outcomes in 146 chest masculinization surgeries.^15^ We showed that patients receiving exogenous testosterone therapy had worse scarring outcomes after chest masculinization surgery, with a testosterone dose– response effect.

Our team also previously developed a model of the GAS hormonal milieu and studied the effects of exogenous testosterone on cutaneous wound healing (WH) in mice, finding that T significantly impairs WH.^11^ However, there are significant limitations in small animal models to capture translationally relevant differences in scarring due to well-established histological skin differences between rodents and humans.^16^ We therefore sought to develop a GAS hormonal milieu model in swine, a more robust option for modeling human scarring, and use this model to test our hypothesis that exogenous testosterone therapy alters scarring.

## Methods

### Swine model

A total of six Hanford adult (6-month-old) miniature swine were included in the study (Sinclair BioResources, Auxvasse, MO, USA). A central venous indwelling catheter was surgically inserted into each swine under general anesthesia 14 days before cutaneous wounding. XX swine (n=4) received bilateral ovariectomies (OVX) while XY swine (n=2) were castrated.

Animals underwent dorsal cutaneous wounding and hormone induction on POD0, two weeks after initial surgery. Excisional 3.5cm wounds (n=10) were created with a #15-surgical blade on the dorsum of each swine without excising fascia or muscle (**Fig. 1**). Wounds were dressed individually with Tegaderm (3M, Saint Paul, MN, USA) and subsequently collectively dressed with gauze pads, Ioban Antimicrobial Incise Drape (3M, Saint Paul, MN, USA), and Vetrap elastic bandages (3M, Saint Paul, MN, USA). Wounds were cleaned and dressings were changed every 72 hours for the first two weeks and every 5 days thereafter.

**Figure 1.**
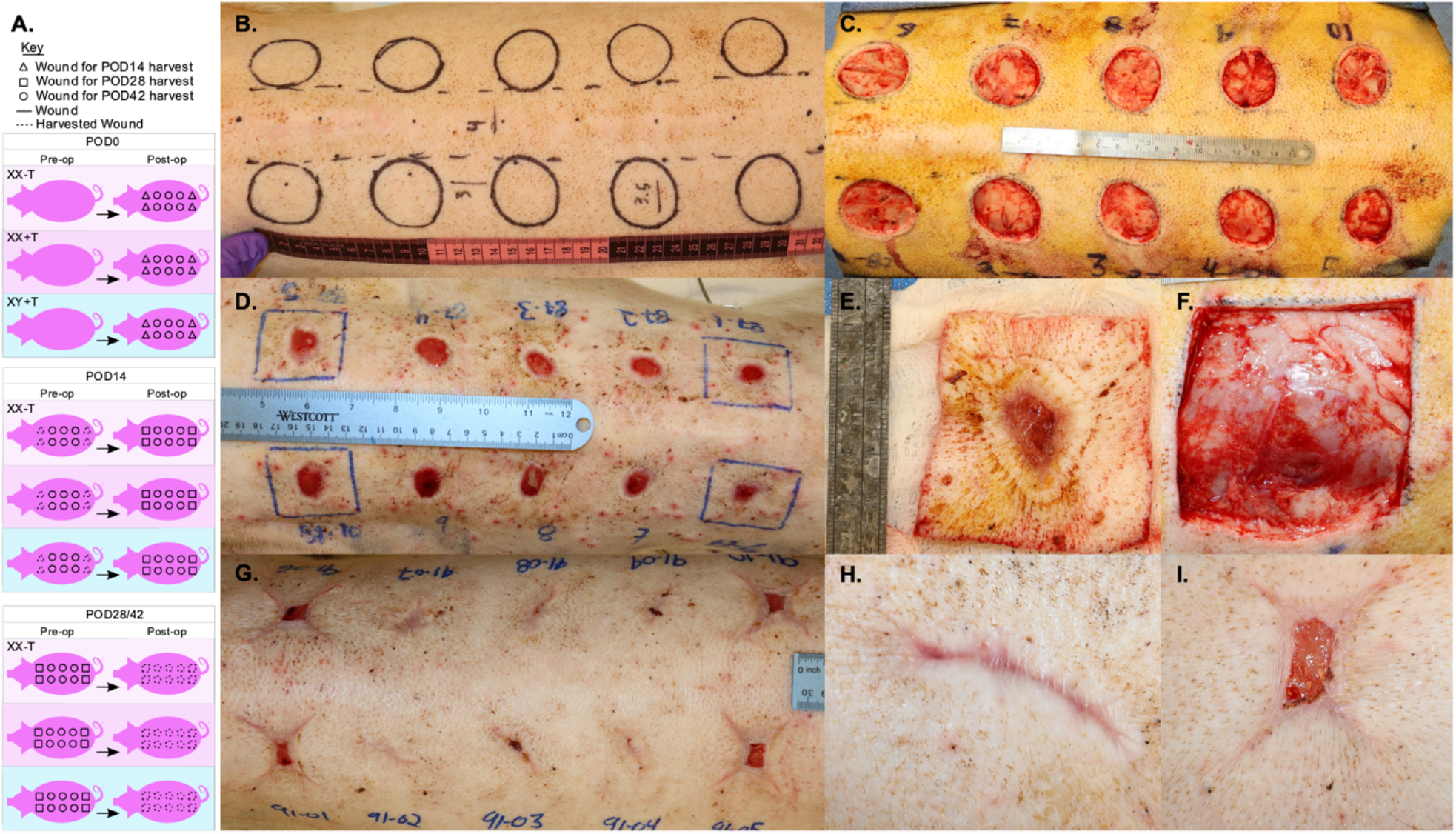
Swine cutaneous scarring model. (**A**) Six swine underwent castration (ovariectomy/orchiectomy) and were randomly assigned to no testosterone or biweekly testosterone therapy. (**B**,**C**) Ten 3.5 cm dorsal excisional wounds were created on each pig. (**D, E, F**) To mimic a chronic wound, four wounds on each swine were re-excised at POD14. Six weeks after initial wounding (**G**), POD42 scar tissue (**H**) and POD28 chronic wounds (**I**) were harvested six weeks after initial cutaneous wounding.

Four wounds from each swine were re-excised on POD14 with a #15-surgical blade to reset the wound bed and mimic delayed chronic wound healing. Six weeks after the initial wounding, scars (POD42) and chronic wounds (POD28) were harvested (**Fig. 1**). All protocols were approved by the Johns Hopkins Animal Care and Use Committee.

### Hormone dosing and measurement

The testosterone regimen was based on our lab’s previous small animal models as well as work conducted by Kothmann et al.^11,17^ Testosterone cypionate (West-Ward Pharmaceuticals Eatontown, NJ, USA) 200mg/ml was diluted with pharmaceutical-grade sesame oil (Sigma-Aldrich, St. Louis, MO, USA) to reach a final concentration of 100mg/ml. Four swine (two XX and two XY) received biweekly 1mg/kg/dose testosterone injections while the remainder received placebo sesame oil.

Testosterone and estradiol levels were measured both in serum and scar tissue samples. Blood samples were collected twice a week and centrifuged to separate serum. Serum was stored at - 20°C and sex hormone levels were analyzed via liquid chromatography-mass spectrometry (LC-MS) (Texas A&M University, College Station, TX, USA). Tissue samples of around 200mg were homogenized in hexane saturated with methanol. The collected extract was evaporated to dryness and reconstituted with acidic aqueous solution. Standard liquid chromatography-mass spectrometry methods for sample preparation and determination of steroid levels were subsequently followed (BRAC Lab, Brigham & Women’s Hospital, Boston, MA, USA).

### Tensile strength testing

A strip of scar tissue was harvested from the same location in each healed wound. Dimensions of excised scars were measured to normalize by cross-sectional area (i.e., scar thickness). Strips of excised scar were loaded onto a 100N load cell force-displacement apparatus. Samples were stretched at a constant strain rate (1mm/sec) until failure. Durometer, stress-strain curves, Young’s Modulus (YM), and ultimate tensile burst strength were analyzed. Wilcoxon Mann-Whitney tests were performed and p-value<0.05 were considered statistically significant.

### Histology

Sections of scar and wound tissue were fixed with 10% formalin for 48 hours and embedded in paraffin blocks. Five-micrometer sections of tissue were created and stained with H&E, Masson’s Trichrome, and Herovici Stains (Abcam, Cambridge, MA). Slides were scanned with a Vectra Polaris (Akoya, Marlborough, MA) and a NanoZoomer-SQ Digital slide scanner (Hamamatsu, Shizuoka, Japan). The granulation tissue area and fibrosis tissue area on each slide were measured by three blinded reviewers using Concentriq interphase (Proscia, Philadelphia, USA). Analysis of variance was performed with significance set at <0.05.

The presence of type I collagen and type III collagen in POD42 scar histology slides stained with Herovici was quantified using FIJI software.^18^ Color deconvolution was applied to isolate type I collagen (red) and type III collagen (blue) staining components. Auto-thresholding using the IsoData algorithm was performed to segment type I (165-255) and type III collagen (15-255) across all samples. Area fraction measurements were performed to quantify each collagen type. Statistical analysis was done using One-way ANOVA and GraphPad Prism Software (Boston, MA, USA).

Histology sections were scored in a blinded fashion by a board-certified dermatopathologist for granulation tissue presence, dermal collagen deposition and orientation, and inflammation. We followed a modified scale previously published by Li et al. and Gupta et al.^19,20^ The presence of granulation tissue was scored 1-4 (1: profound, 2: moderate, 3: scanty, 4: absent). Inflammation was scored 1-3 (1: plenty, 2: moderate, 3: few). Collagen fiber orientation was scored 1-3 (1: vertical, 2: mixed, 3: horizontal). The amount of early collagen (type III) and late collagen (type I) were graded 1-3 (1: profound, 2: moderate, 3: minimal). Higher scores indicate better healing and scarring.^20^

### RNA seq

Scar tissue samples (POD42) were homogenized using a GentleMacs tissue homogenizer (Miltenyi Biotec, Gaithersburg, MD). RNA was isolated from the supernatant and quantified using the NanoDrop 2000 spectrophotometer (Thermo Scientific, Wilmington, DE, United States), according to the manufacturer’s protocol. A total of 39 samples underwent total RNAseq using an S2 flow cell. We used Ensembl’s S. scrofa 11.1 release (http://mart.ensembl.org/Sus_scrofa/Info/Index). Counts were quantified at the transcript level and quality control was performed. Differential gene analysis was performed using DESeq2. Gene ontology analysis was performed using TopGO. Pathway analysis was performed by using GSEA (Broad Institute, Boston, MA) using MSigDB 2022 Hallmark and Reactome pathway sets. Additional information is available in supplemental methods.

## Results

### Validation of a clinically relevant porcine testosterone dosing regimen

Serum testosterone levels were consistent over time. Swine injected with exogenous testosterone therapy showed a significant increase in serum testosterone levels relative to the no-T group (5.9ng/dL XXnoT, 189.9ng/dL XX+T, 221.8ng/dL XY+T; p<0.01). Swine serum estradiol levels did not differ between hormone regimen experimental groups (5.07pg/ml XXnoT, 13.48pg/ml XX+T, 8.61pg/ml XY+T; TTest XXnoT vs XX+T p=ns; TTest XXnoT vs XY+T p=ns) (**Fig. 2)**.

**Figure 2.**
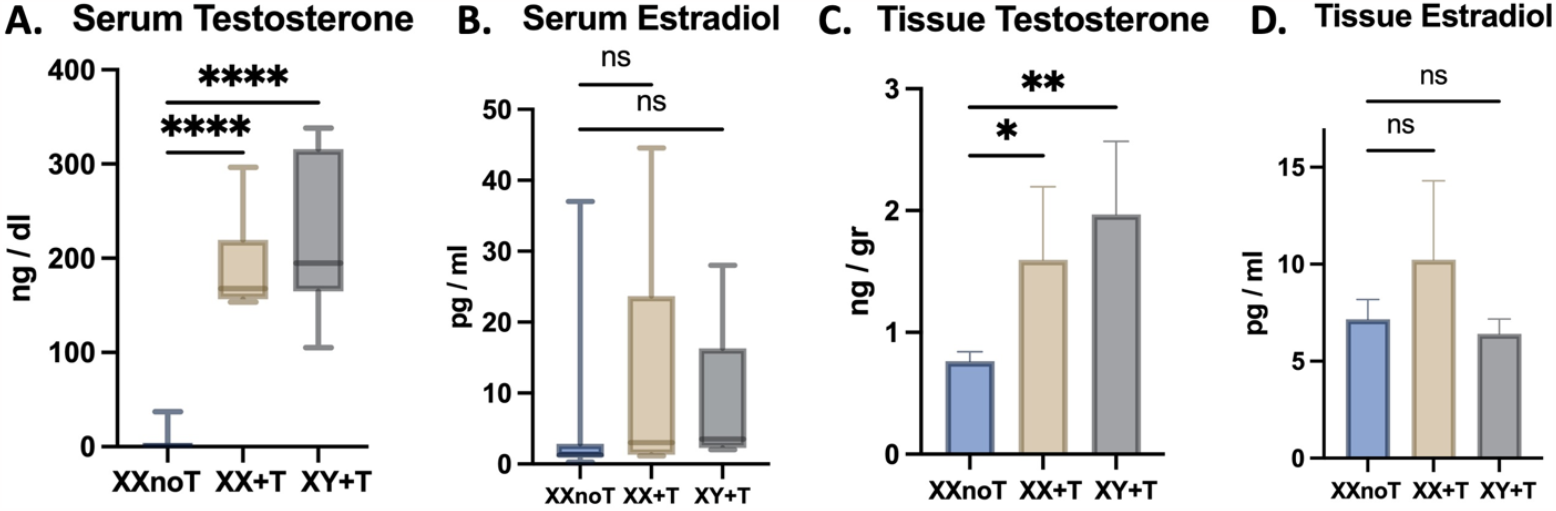
Systemic testosterone therapy yielded physiologic levels in serum and penetrated scar tissue without elevating estradiol levels. (**A**) +T swine showed a significant increase in serum testosterone levels relative to the no-T group. (**B)** Swine serum estradiol levels did not differ between hormone regimen experimental groups. (**C**) Swine scar tissue (POD42) and chronic wound samples (POD28) from animals treated with T showed increased tissue levels of testosterone compared to animals not treated with T. (**D**) Testosterone therapy did not alter serum or tissue estradiol levels significantly.

### Systemic testosterone therapy penetrates scar tissue

Swine scar tissue (POD42) and chronic wound tissue samples (POD28) from animals treated with T showed increased levels of testosterone compared to animals not treated with T (0.76ng/g of tissue XXnoT, 1.60ng/g XX+T, 1.97ng/g XY+T; p=0.007). Estradiol tissue levels did not differ between groups (7.17pg/ml XXnoT, 10.23pg/ml XX+T, 6.39pg/ml XY+T; TTest XXnoT vs XX+T p=0.492, TTest XXnoT vs XY+T p=0.567)

### Testosterone-exposed scars and chronic wounds demonstrate greater histologic thickness, width, and collagen deposition

Histologic analysis of chronic POD28 wounds showed increased granulation tissue (GT) area (0.52cm^2^ for XXnoT, 0.66cm^2^ for XX+T, 0.88cm^2^ for XY+T, p=0.039) as well as increased epithelial thickness in the testosterone-treated groups (0.45mm for XXnoT, 0.62mm for XX+T, 0.68mm for XY+T, p=0.05). POD42 samples from the +T swine showed increased mean scar area compared to XXnoT swine (0.157cm^2^ XXnoT, 0.290cm^2^ XX+T, 0.268 cm^2^ XY+T; p=0.007). Mean scar thickness was also increased in the scars from T-treated swine (0.246mm XXnoT, 0.389mm XX+T; 0.423mm XY+T; p<0.001) (**Fig. 3**).

**Figure 3.**
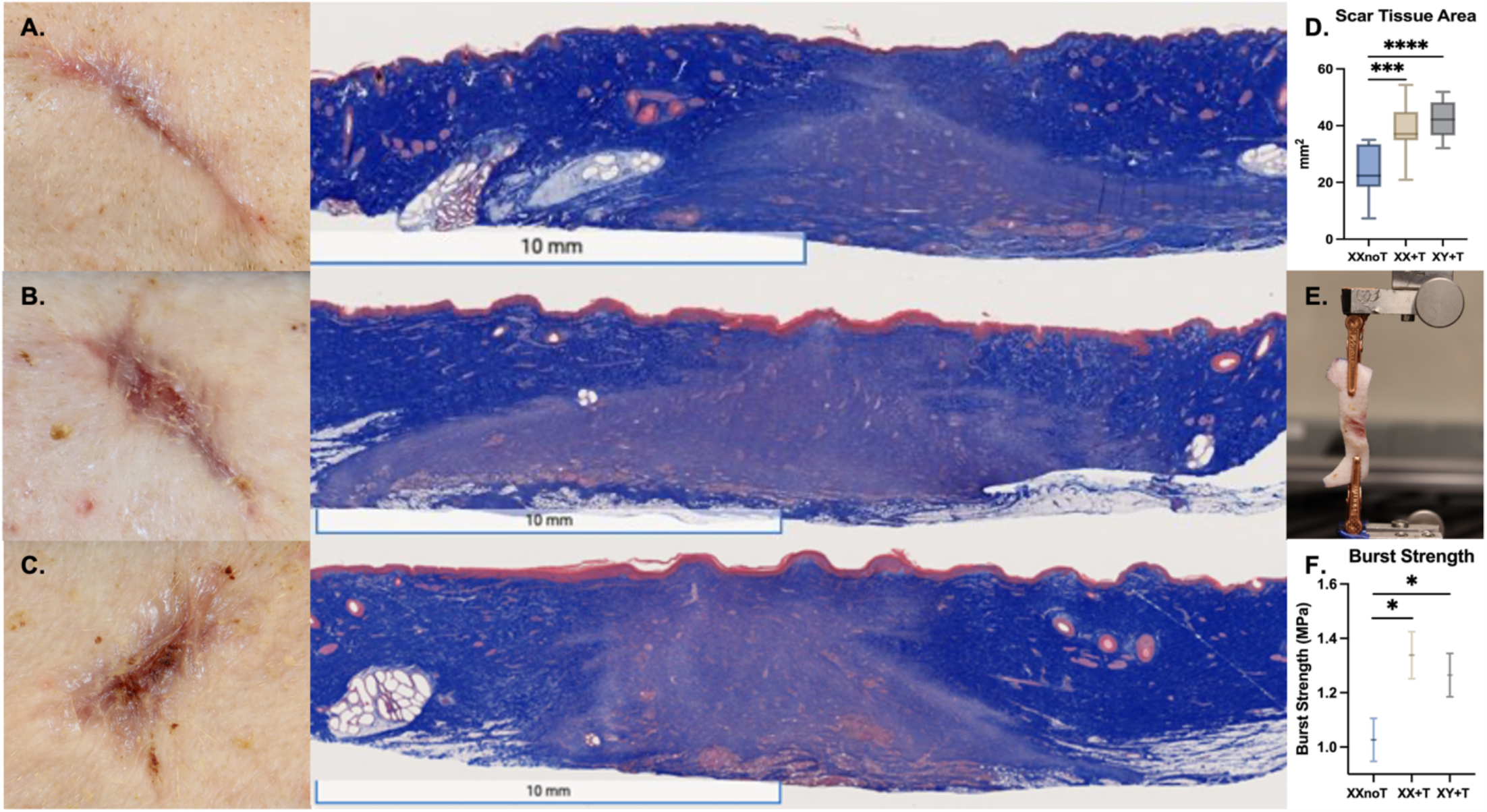
Testosterone-exposed scars demonstrate greater scar tissue area (mm^2^), but also a higher CSA-adjusted burst strength per mm^2^ of scar. POD42 scars from (**A**) XXnoT, (**B**) XX+T, and (**C**) XY+T swine. (**D**) POD42 scars from the +T swine showed increased mean scar tissue area compared to XXnoT swine. **(E)** Tensile strength was tested using a 100N load cell force-displacement apparatus. Samples were stretched at a constant strain rate (1mm/sec) until failure. (**F**) Cross-sectional area-adjusted burst strength of scars from swine exposed to T were higher compared to controls (0.81 MPa for all -T vs 1.30 MPa for all +T; p=0.001, 0.81 XXnoT vs 1.29 MPa XX+T p=0.02; 0.81 MPa XXnoT vs 1.31 MPa XY+T p<0.01).

Color deconvolution analysis of histologic scar tissue stained with Herovici showed testosterone was associated with an increase in type I collagen deposition (24.69 XXnoT, 62.83 XX+T, 62.39 XY+T; p<0.05) as well as type III collagen in the XX+T group compared to control (59.49 XXnoT, 75.90 XX+T, 62.01 XY+T; p<0.05) (**Fig. 4**). T-exposed scars were associated with an increase in type I to III collagen ratio compared to controls (0.46 XXnoT, 0.81 XX+T, 1.03 XY+T; p<0.05) (**Fig. 4**).

**Figure 4.**
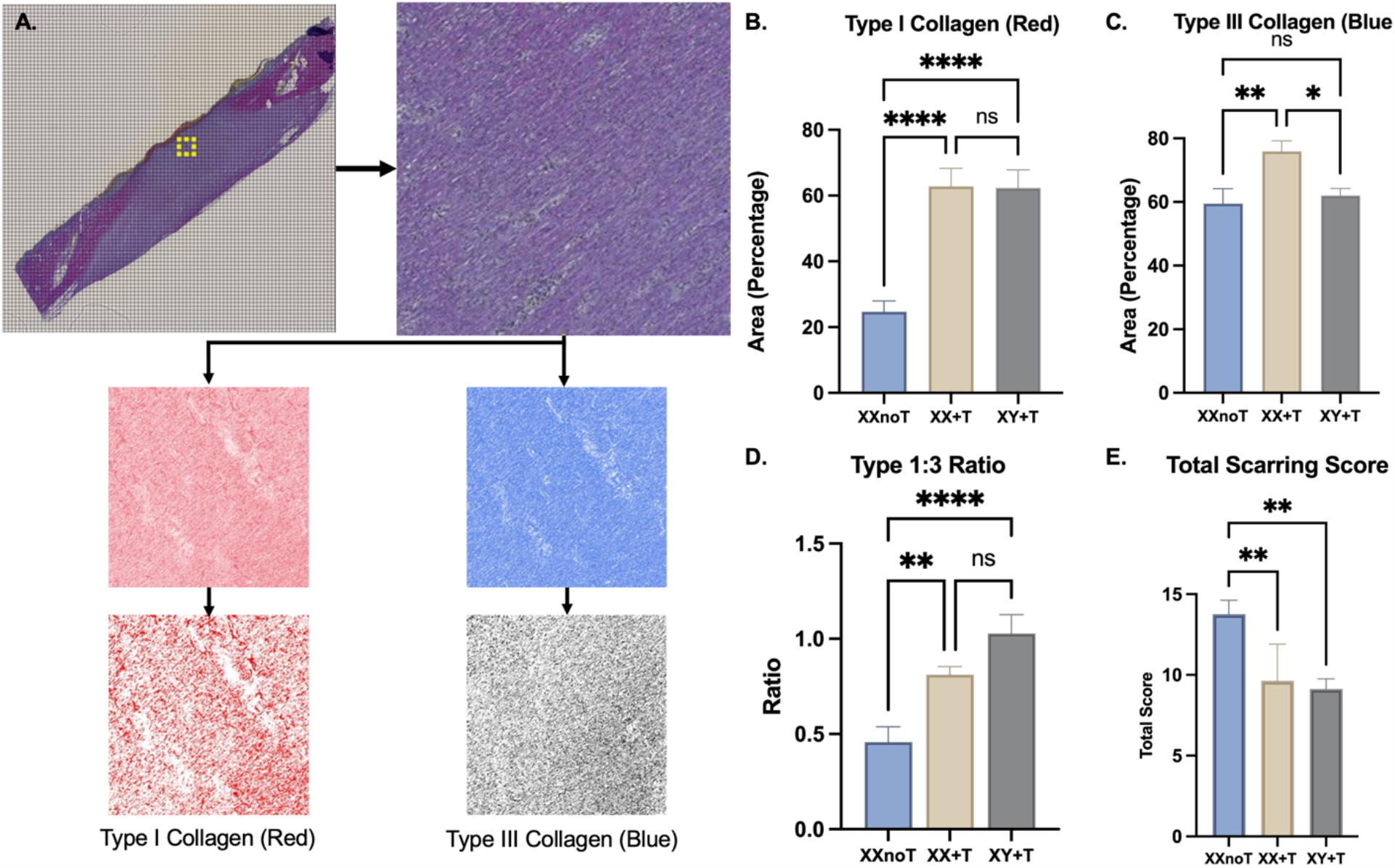
Testosterone-exposed scars show greater deposition of both type I and III collagen, increased type I to III collagen ratio, and worse overall scarring scores. (**A**) Color deconvolution analysis of Herovici-stained POD42 scars showed testosterone-exposed scars had (**B**) greater deposition of type I and (**C**) type III collagen as well as (**D**) higher type I to type III ratio. (**E**) Dermatopathologist assessment using histologic scoring scales found testosterone led to significantly more pathologic tissue repair.

### Dermatopathologist assessment showed testosterone induced more pathologic tissue repair

Scars exposed to exogenous testosterone had worse overall average scarring scores compared to controls (13.75 XXnoT, 9.63 XX+T, 9.13 XY+T; p<0.05). Dermatopathologist scores showed T-treated scars had a decrease in granulation tissue and inflammation present as well as an increase in vertical collagen fiber orientation, commonly associated with poor wound healing and scarring (p>0.05) (**Fig. 4**).

### Testosterone-exposed scars develop greater tensile strength

The mean and standard deviation of YM of 42-day (0.01000 MPa, 0.00743 MPa) scars was slightly higher than 28-day (0.00738 MPa, 0.00548 MPa) wounds, but not significantly so. Adjusted burst strength of scars from swine exposed to T were higher compared to controls (0.81 MPa for all -T vs 1.30 MPa for all +T; p=0.001, 0.81 XXnoT vs 1.29 MPa XX+T p=0.02; 0.81 MPa XXnoT vs 1.31 MPa XY+T p=<0.01). There was no significant difference in burst strength between XX+T and XY+T. **(Fig. 3)**.

### Testosterone-exposed scars show upregulation of metabolic, ECM and immune gene sets

Gene ontology enrichment analysis was analyzed by cellular component, molecular function, and biological processes. Gene sets related to metabolism, immune response to infection, and T-cell/leukocyte response were the most differentially expressed in +T scars. Gene set enrichment analysis (GSEA) showed that Reactome pathways related to laminin interactions, keratinization, ECM turnover and NCAM1 interactions were upregulated in +T scars. Fewer pathways were downregulated (**Fig. 5**). Hallmark pathway analysis showed upregulation in gene sets associated with myogenesis and hedgehog signaling in +T scars, with downregulation of interferon and anti-viral defense pathways. (**Fig. 5**).

**Figure 5.**
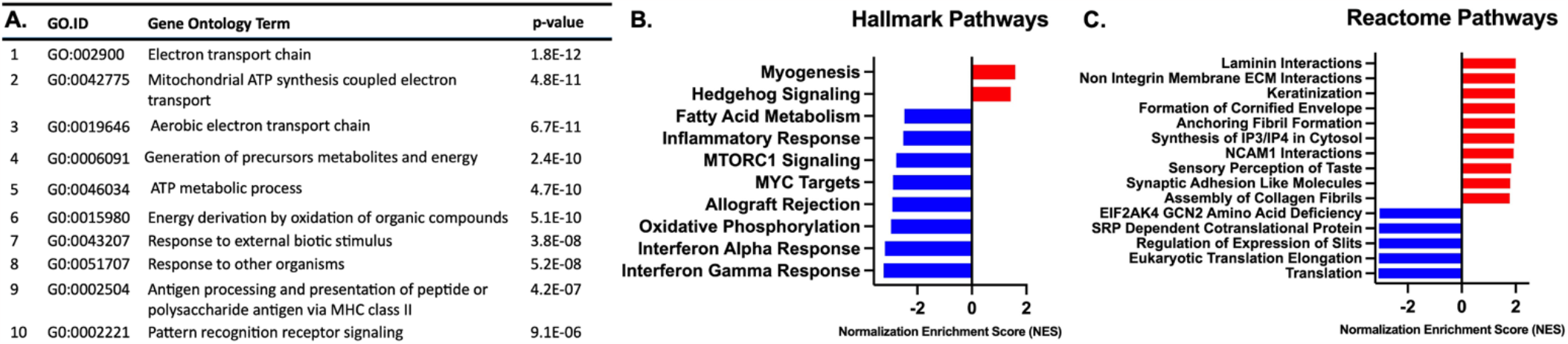
Testosterone induces significant shifts in metabolic, ECM and immune gene pathways during tissue remodeling. (**A**) Gene ontology analysis showed testosterone exposure was associated with significant alterations in cellular metabolism and immune response gene sets. (**B**) Hallmark pathway analysis showed testosterone exposure was associated with an upregulation in gene sets related to myogenesis, but strong downregulation of interferon-mediated immune and catabolic pathways. (**C**) Reactome pathway analysis showed testosterone was associated with upregulation of pathways related to keratinization and formation of collagen and laminin.

## Discussion

We developed a GAS hormonal milieu model in swine to investigate how exogenous testosterone therapy alters scarring. First, we created and validated the preclinical porcine model of exogenous testosterone therapy. We hypothesized that tissue hormone levels would be related to serum hormone levels and developed a protocol to test this. Given that estrogen has been proven to improve wound healing, we tested estradiol levels from both serum and scar samples and found no differences between group 17-b-estradiol levels in both serum and tissue.^21–23^ We demonstrated that systemic testosterone therapy does penetrate scar tissue (i.e., +T swine have higher scar tissue testosterone vs. noT), making testosterone-induced effects on tissue remodeling physiologically plausible.

We demonstrated that +T swine showed increased mean scar area and thickness compared to noT swine. Chronic wounds also showed increased granulation tissue area in +T compared to noT swine. Color deconvolution analysis of Herovici-stained POD42 scar slides and dermatopathologist scores showed testosterone-exposed scars were associated with a decrease in late granulation tissue and inflammatory infiltrate as well as an increase in collagen I deposition and type I vs. type III collagen ratio. These findings complement previous literature which associate an increased overall collagen deposition and higher ratio of type I to type III collagen in the pathogenesis of hypertrophic scarring and keloid formation.^24–26^

This study is also the first to assess how testosterone impacts the tensile strength and elasticity of pig scars. The observed increase in tensile burst strength of scars from both XX+T and XY+T swine compared to noT swine suggests that exogenous testosterone has a strong influence on the mechanical properties of scars regardless of chromosomal sex. This is unsurprising as a broad variety of literature has examined gonadal hormone vs chromosomal effects, and the latter tend to be much more subtle.^27–31^ Generally, the phenotypic effects of testosterone were observed to be broadly similar in all measurements between the XY+T and XX+T pigs.

Testosterone appears to increase the physical strength of scars, possibly via supraphysiologic deposition of collagen and other extracellular matrix factors. We hypothesize that testosterone induces early excess angiogenesis and proliferation of granulation tissue which may then lead to increased scar tissue development. Scar formation and remodeling typically begin two to three weeks after injury occurs and can last over a year. In remodeling, type III collagen and proteoglycans are replaced with type I collagen, and the orientation of collagen fibrils becomes more organized. This collagen rearrangement corresponds with an increase in wound tensile strength. Unlike collagen, elastin is absent from the granulation tissue deposited by fibroblasts. The lack of elastin, which provides elastic recoil for the dermal matrix, results in a more rigid and inelastic extracellular matrix.

Reactome and hallmark pathway analysis revealed testosterone was associated with an upregulation of pathways related to keratinization, myogenesis, hedgehog signaling, and formation of collagen and laminin, congruent with previous findings. Gene set analysis showed an upregulation of cellular metabolism as would be expected by testosterone’s pro-glycolytic effects, but also elucidated changes in immune response gene sets, suggesting T effects on immune cells may be a mechanism for altering scar formation.

Traditional practices of stopping testosterone perioperatively were based on concerns about thrombocytosis, hypertension and thromboembolic events, but recent studies suggest these concerns are largely theoretical.^32^ Because hormone therapy provides TGD patients with psychological benefits, patients may experience distress if asked to pause an important aspect of gender-affirming care, and cessation is less common now.^33^ However, these newer studies do not address scar results, potentially resulting in perioperative use of a medication that delays WH and increases scar hypertrophy. Our findings suggest that continued use of exogenous testosterone during the perioperative period could bring greater scarring morbidity.^34^ It is therefore important to find therapeutic approaches that exert localized anti-androgenic properties without systemic effects. The development of topical antiandrogens that competitively bind to androgen receptors represents a promising area of future research as a novel approach to controlling the tissue repair cascade through the sex hormone axis. ^32^

## Conclusions

We developed a novel preclinical model to study the effects of the sex hormone testosterone on scarring. Testosterone induces early proliferation of excessive granulation tissue, which eventually leads to increased scar tissue. Testosterone appears to increase the physical strength of scars via supraphysiologic deposition of collagen and other ECM factors.

## Acknowledgments

We thank the American Association of Plastic Surgeons (AAPS) for partially funding this project. Dr. Coon was the 2021 recipient of the Furnas Academic Scholar Award.

## List of abbreviations

noT: No testosterone experimental group
+T: Experimental group exogenously treated with biweekly testosterone cypionate
POD42: Postoperative day 42. Scar tissue excised 42 days after initial wounding
POD28: Postoperative day 28. Scar tissue excised 28 days after partial harvest
TGD: transgender and gender nonbinary individuals
GAC: gender-affirming care
GAS: Gender-affirming surgery
T: Testosterone
WH: Cutaneous Wound Healing
OVX: Bilateral ovariectomies
LC-MS: liquid chromatography-mass spectrometry
YM: Young’s Modulus
GSEA: Gene set enrichment analysis

## Supplemental Methods

FASTQ-format files were submitted to FastQC (v0.11.9) to check for basic sequence statistics and identify poor quality samples. Following this, the FASTQ files were aligned to the Ensemble S. Scrofa v11.1 reference using the STAR (v2.7.10a aligner). Output BAM-format files were sorted using samtools (v1.16.1) and duplicate reads were identified with Picard Tools MarkDuplicates (v2.27.4). We retained these duplicate reads for downstream differential expression, but three samples with excessive duplication and low alignment rates were removed from analysis. Raw read counts were quantified using the SubRead featureCounts (v2.0.3) package. Differential expression analysis was conducted using DESeq2 (v1.34.0).

## Notes

**Data Availability:** The data that support the findings of this study are available from the corresponding author upon reasonable request.

**Conflicts of Interest:** The authors do not have any pertinent conflicts of interest to disclose.

### Competing Interest Statement

The authors have declared no competing interest.

### Summary of Updates

The figures were improved.

